# Sperm meet the elevated energy demands to attain fertilization competence by increasing flux through aldolase

**DOI:** 10.1101/2025.04.09.647926

**Authors:** Sara Violante, Aye Kyaw, Lana Kouatli, Kaushik Paladugu, Lauren Apostolakis, Macy Jenks, Amy Johnson, Ryan D. Sheldon, Anthony L. Schilmiller, Pablo E Visconti, Justin R Cross, Lonny R. Levin, Jochen Buck, Melanie Balbach

## Abstract

Prior to ejaculation, sperm are stored in the epididymis in a ‘resting’ metabolic state. Upon ejaculation, sperm must alter their metabolism to generate the energy needed to support the motility and maturation process known as capacitation to reach and fertilize the oocyte. How sperm regulate the capacitation-induced increase in carbon flux is unknown. Here, we use ^13^C stable isotope labeling to follow glucose metabolism through sperm central carbon metabolic network before and after sperm activation. We identify regulatory steps which sperm use to alter their metabolic state from resting to highly active. In activated sperm, glucose flux through glycolysis is increased at the expense of the pentose phosphate pathway to increase energy yield. Increased glycolytic activity seems to be due to capacitation-induced stimulation of flux through aldolase. In the mitochondria-containing midpiece, glycolytically generated pyruvate feeds the TCA cycle to further maximize energy yield via oxidative phosphorylation. In the mitochondria-free principal piece of the tail, pyruvate produced from glycolysis is reduced to lactate by lactate dehydrogenase. Reduction to lactate regenerates oxidized NAD^+^ ensuring a sufficient supply to support glycolysis. The resultant lactate is at least partially secreted. Finally, we find evidence that there is an as yet unknown endogenous source of energy in sperm feeding the upregulation of TCA cycle intermediates. These studies provide the most complete picture of the metabolic shift which occurs in capacitating sperm.

**Significance statement:** A rapid switch from a quiescent to a high energy-demanding state during ejaculation is essential for sperm to reach and fertilize the oocyte. Somatic cells also undergo bioenergetic switches from low to very high energy demand. However, because metabolic processes essential for proliferation are going on in parallel, it is difficult to identify the molecular mechanisms regulating the increase in ATP production. This study represents the first complete picture of the metabolic reprogramming that happens in sperm upon ejaculation. Using stable isotope labeling, we identify rate-limiting enzymatic steps and points of regulation directing the changes in metabolic flux. Our sperm metabolic studies allow us to identify conserved mechanisms of metabolic regulation that are crucial for the survival of mammalian cells.

## Introduction

Sperm are highly specialized cells designed to fertilize the oocyte and deliver the haploid male genome. In mammals, sperm are produced in the testis and attain the capability for motility as they pass through the epididymis. Upon reaching the cauda of the epididymis where they are stored awaiting ejaculation, morphologically mature and transcriptionally and translational silent sperm are at rest, with relatively low energy demand. With ejaculation, sperm have to rapidly enhance their metabolism^1,2^ to produce the energy needed to support flagellar movements propelling them during their journey to the oocyte, to acquire fertilization competence (capacitation^3,4^), and to successfully fertilize the oocyte.

Studying metabolism in mouse sperm affords unique opportunities to examine the metabolic reprogramming from physiologically dormant to fully active states. Starting from sperm surgically extracted from the cauda epididymis, *in vitro* incubation in capacitating media triggers sperm movement, initiates capacitation, and induces the metabolic profile resembling ejaculated mouse and human sperm^5^. By directly comparing epididymal mouse sperm incubated in capacitating versus non-capacitating media, we demonstrated that capacitating sperm increase glucose consumption^6^ and elevate rates of glycolysis and oxidative phosphorylation (oxphos)^7,8^.

The molecular changes of capacitation are initiated by a signaling cascade driven by the bicarbonate in seminal fluid, which is mimicked in capacitation media, resulting in activation of soluble adenylyl cyclase (sAC; Adcy10)^9–11^ and protein kinase A (PKA)^12,13^. This bicarbonate-dependent sAC/PKA signaling cascade leads to phosphorylation of ion channels, metabolic enzymes, and structural proteins^14–16^ which mediate the changes essential for sperm to fertilize the oocyte. Among these changes, during capacitation sperm switch to a high-amplitude, asymmetric, hyperactivated flagellar beating pattern which allows sperm to generate the propulsive forces needed to reach the oocyte and penetrate its vestments^17^. Because sperm from sAC null mice, as well as wild type sperm in the presence of sAC inhibitors, do not undergo capacitation^18–22^, mice also provide the tools necessary to identify whether observed metabolic changes are dependent upon capacitation-induced intracellular signaling. For example, the previously reported increase in glucose consumption during capacitation is dependent upon cAMP signaling^6^, confirming it is a consequence of capacitation-induced signal transduction.

Like somatic cells, mammalian sperm possess the metabolic machinery to generate ATP and other high-energy compounds via glycolysis, pentose phosphate pathway (PPP), and oxphos^23–26^. However, unlike somatic cells, sperm metabolism is focused exclusively on energy production. After delivering the pronucleus to the oocyte, the body of the sperm is discarded. In somatic cells, the PPP produces NADPH facilitating reducing reactions and ribose-5-phosphate to synthesize nucleic acids, and mitochondria are hubs for generating amino acids, free fatty acids, cholesterol, and other biomolecules essential for cellular growth. In contrast, sperm metabolism is not focused on generating biomass. Also distinct from other mammalian cells, it is proposed that sperm’s unique architecture separates their metabolic machinery. The sperm tail is organized into two separate sections, midpiece and principal piece. Glycolytic enzymes are most abundantly found in the principal piece of the sperm tail, which is devoid of mitochondria. The sperm’s mitochondria with TCA cycle and oxphos machinery are located exclusively in the midpiece. This unique compartmentalization fuels an ongoing debate regarding which pathways are used during the sperm’s journey to the oocyte and whether glycolysis and oxphos are connected as in somatic cells^27–33^. We previously found that capacitation increases glucose uptake in both midpiece and principal piece^7^. Using extracellular flux analysis we showed that the capacitation-induced increase in oxphos is dependent on glycolysis^7^ and NMR studies using U-^13^C-glucose revealed that exogenous glucose can be converted into the TCA metabolite citrate^34^. Combined, these studies suggest that glycolytically-generated substrates can directly fuel mitochondrial oxphos.

Despite these advances, it remains unknown how capacitation contributes to metabolic reprogramming in mammalian sperm. To investigate how individual endogenous metabolites change following exposure to capacitating conditions, we developed mass spectrometry and metabolic profiling methodologies for mammalian sperm^8^. Unsupervised hierarchical clustering revealed distinct metabolite patterns for epididymal mouse sperm incubated in non-capacitating and capacitating conditions^8^. Specifically, capacitation-induced changes in the rates of glycolysis and oxphos detected via mass spectrometry were consistent with extracellular flux analysis^7^: Mouse sperm incubated in capacitating conditions showed increased levels of lactate and citrate, suggesting a capacitation-induced increase in carbon flux through glycolysis and the TCA cycle^8^. Metabolic profiling provides information on the abundance of each metabolite. Analysis of metabolite pool sizes has limitations as it cannot identify reactions operating at increased or decreased rates, and, therefore, fails to distinguish between metabolite build-up due to increased production, or decreased consumption. In contrast, stable isotope labeling (SIL) analysis reveals metabolite production and consumption kinetics^35^. Here, we follow the changes in the enzyme-regulated steps composing the sperm central carbon metabolic network during capacitation using SIL with ^13^C-isotopically labeled glucose through metabolites of glycolysis, the PPP, and the TCA cycle. We fed U-^13^C-labeled glucose to non-capacitating or capacitating epididymal mouse sperm and analyzed labeling ratios of downstream metabolites by mass spectrometry at distinct time points. By following the distribution of labeled carbon atoms over the metabolic network we identified rate-limiting enzymatic steps and points of regulation directing the changes in metabolic flux.

## Results

### Metabolic flux in non-capacitating and capacitating sperm

To decipher the dynamic changes in sperm central carbon metabolism during capacitation, we applied SIL analysis to non-activated and activated mammalian sperm. The sensitivity of mass spectrometry combined with the robustness of ion-pairing chromatography allowed us, for the first time for male germ cells, to simultaneously follow the flux of glucose-derived heavy carbon through glycolysis, the TCA cycle, and the PPP. *Mus musculus* C57Bl/6 mouse sperm were isolated from the cauda epididymis into media containing minimal glucose concentrations (0.56 mM) to avoid the induction of starvation pathways^36,37^ and were mixed with either non-capacitating sperm buffer (TYH) or conditions that induce molecular hallmarks of capacitation (TYH + 3 mg/ml BSA + 25 mM HCO_3_ ^−^). Sperm were simultaneously subjected to uniformly-labeled 5.6 mM U-^13^C-glucose where all six carbons are replaced by ^13^C (Fig. 1). Not more than 90% labeling can be achieved due to the initial unlabeled U-^12^C-glucose present in media. Metabolism was quenched at distinct time points between 0 and 90 min and relevant metabolites were detected by LC-MS. The incorporation of ^13^C labeling was followed by measuring all isotopologues (m+0 to m+6). The relative labeling percentage (Fig. 1B) and abundance of each isotopologue (Fig. 1C) for each metabolite was determined.

**Fig. 1:**
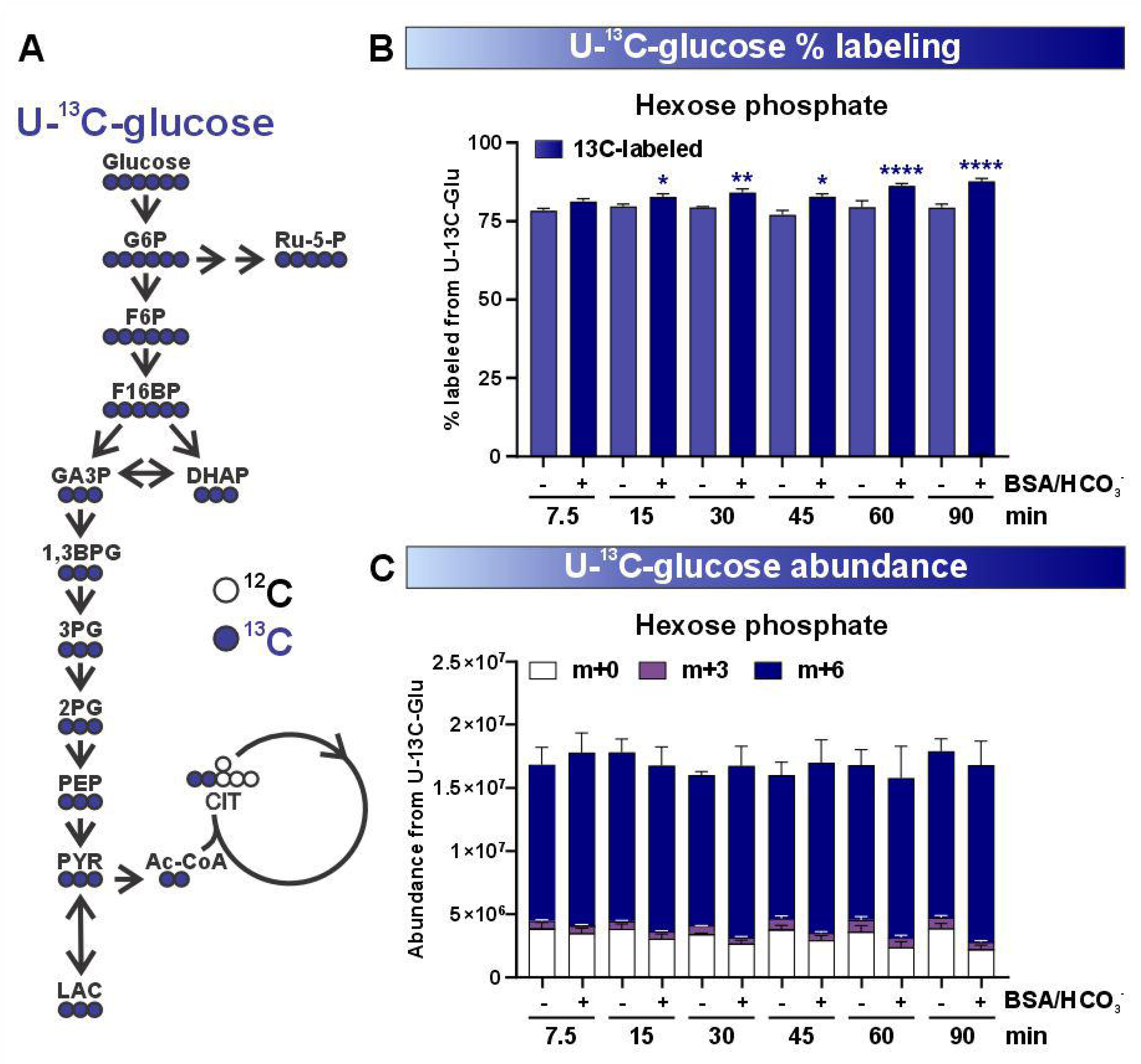
Metabolic flux in non-capacitating and capacitating sperm. **(A)** Schematics of metabolic flux through central carbon metabolism from U-^13^C-glucose. **(B,C) (B)** Fractional enrichment of total labeled (^13^C, m+1 to m+6 isotopologues) and **(C)** total ion intensities of unlabeled (^12^C, m+0) and labeled (^13^C, m+3 or m+6 isotopologues) hexose phosphate in non-capacitating (-BSA/HCO_3_ ^+^) and capacitating (+BSA/HCO_3_ ^+^) epididymal mouse sperm in U-^13^C-glucose. Sperm were incubated in an initial concentration of 0.56 mM of unlabeled U-^12^C-glucose followed by the addition of labeled 5.6 mM U-^13^C-glucose and collected at the indicated time points. Mean + SEM, n≥4. Differences between conditions were analyzed using one-way ANOVA compared to sperm incubated in non-capacitating conditions at 7.5 min (white asterisk: m+0, blue asterisk: m+3/m+6), *P<0.05, **P< 0.01, ***P<0.001, ****P<0.0001.

Incubating epididymal mouse sperm with U-^13^C-glucose revealed a rapid uptake and conversion of U-^13^C-glucose into m+6 hexose phosphate (Fig. 1B). The isomers ^13^C-glucose-6-phosphate and ^13^C-fructose-6-phospate are not distinguished in our chromatographic method and are therefore referred to as hexose phosphate. At the first time point of 7.5 min, there is approximately 75% labeling from U-^13^C-glucose, close to our maximal possible labeling of 90% (Fig. 1B). At 7.5 min labeled hexose phosphate abundance and labeling percentages were not significantly different between non-capacitating and capacitating sperm (Fig. 1B,C); thus, any capacitation-induced changes in amount of labeled glucose-6-phosphate (G6P) and conversion into fructose-6-phosphate (F6P) reflecting the previously observed increased glucose uptake^6,28^ were already equilibrated by this early time point. At later time points, fractional labeling increases in capacitating sperm presumably due to diminution of endogenous U-^12^C-glucose (Fig. 1B,C). As expected from using U-^13^C-glucose, the most abundant isotopologue was m+6; however, the m+3 isotopologue was also detected, suggesting that sperm also convert three-carbon metabolites into hexoses (Fig. 1C).

### Capacitating sperm accumulate and secrete more lactate, confirming increased glycolytic flux

We began our studies of glucose metabolism by asking whether the previously reported intracellular accumulation and release of lactate in mammalian sperm incubated with U-^13^C-glucose^38–42^ was confirmed using our approach (Fig. 2A,B). Indeed, the intracellular accumulation of m+3 lactate was time-dependent and, as observed by NMR^34^, enhanced in capacitating sperm. Capacitation also increased secretion of lactate into the surrounding media (Fig. 2C,D). Among the glycolytic intermediates we could detect, lactate was the only metabolite secreted into sperm media. We detected a time-dependent increase of m+3 lactate in the media of both non-capacitating and capacitating sperm incubated with U-^13^C-glucose (Fig. 2C,D). As with intracellular lactate, lactate secretion was potentiated in capacitation media^34^.These data confirm that capacitating sperm metabolize more glucose into lactate than non-capacitating sperm.

**Fig. 2:**
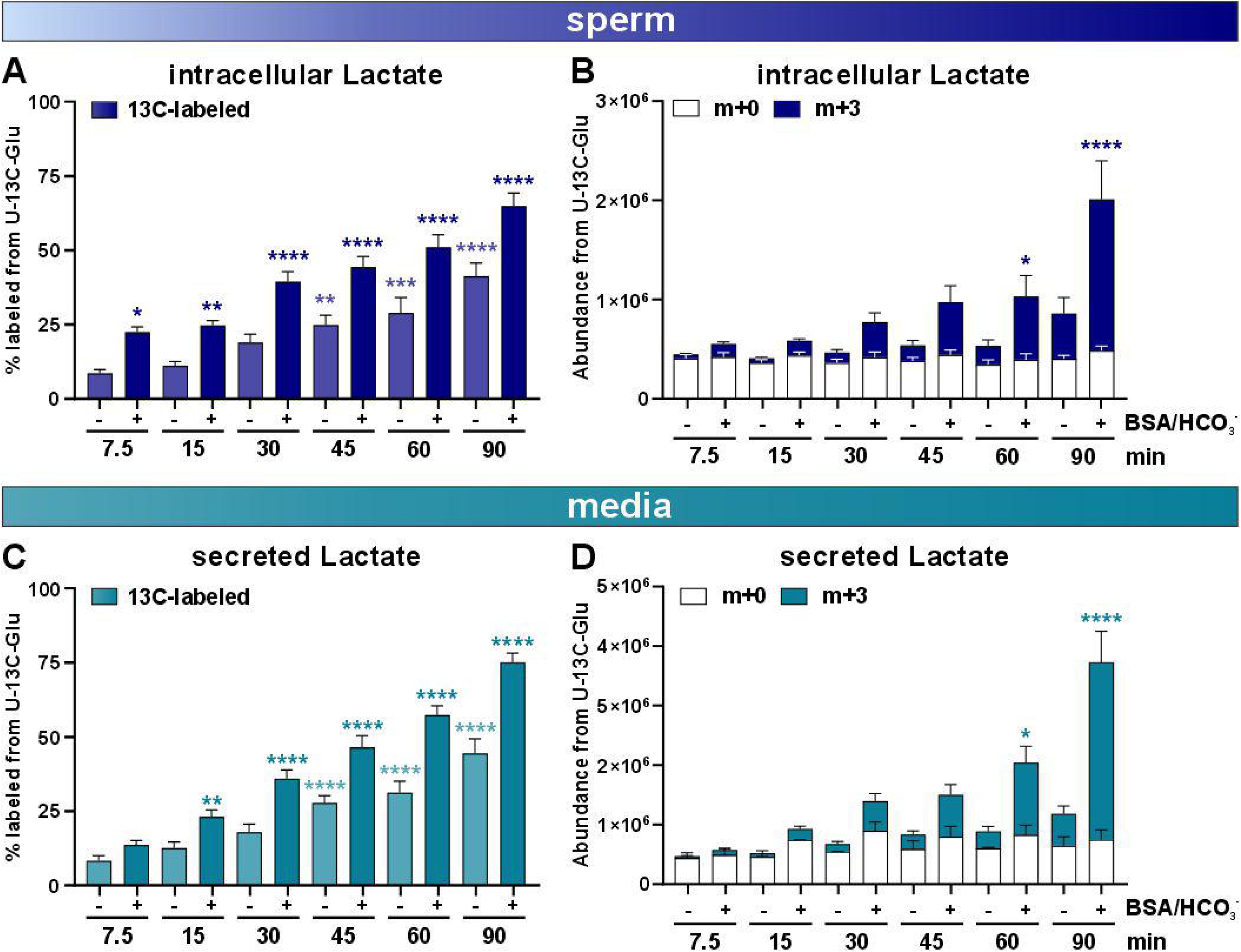
Capacitating sperm accumulate and secrete more lactate. **(A,B) (A)** Fractional enrichment of total labeled (^13^C, m+3 isotopologue) and **(B)** total ion intensities of unlabeled (^12^C, m+0) and labeled (^13^C, m+3 isotopologues) lactate in non-capacitating and capacitating sperm incubated for the indicated time points in 5.6 mM U-^13^C-glucose. Mean + SEM, n≥7. **(C,D) (C)** Fractional enrichment of total labeled (^13^C, m+3 isotopologue) and **(D)** total ion intensities of unlabeled (^12^C, m+0) and labeled (^13^C, m+3 isotopologues) lactate levels in the media surrounding non-capacitating and capacitating sperm incubated for the indicated time points in 5.6 mM U-^13^C-glucose. Mean + SEM, n≥7. Differences between conditions were analyzed using one-way ANOVA compared to sperm incubated in non-capacitating conditions at 7.5 min; (blue/green asterisk: m+3), *P<0.05, **P< 0.01, ****P<0.0001.

sAC-dependent increase of cAMP is the essential initial signal eliciting the molecular changes required for fertilization competence. Therefore, we repeated U-^13^C-glucose SIL experiments with sperm from sAC knockout (sAC KO) mice or sperm from wild type mice where sAC was pharmacologically inhibited by incubation with the sAC inhibitor TDI-11861 during *in vitro* capacitation or *in vivo* (i.e., sperm were isolated from mice pre-treated with sAC inhibitor TDI-11861)^18,19,22^ to determine whether the observed increase in glycolytic activity was due to capacitation-related signaling. In sperm devoid of sAC, the capacitation-induced increase in m+3 lactate accumulating in sperm and secreted into the media was absent (Fig. 3); therefore, the increased flux through glycolysis into lactate is a consequence of capacitation-induced signaling.

**Fig. 3:**
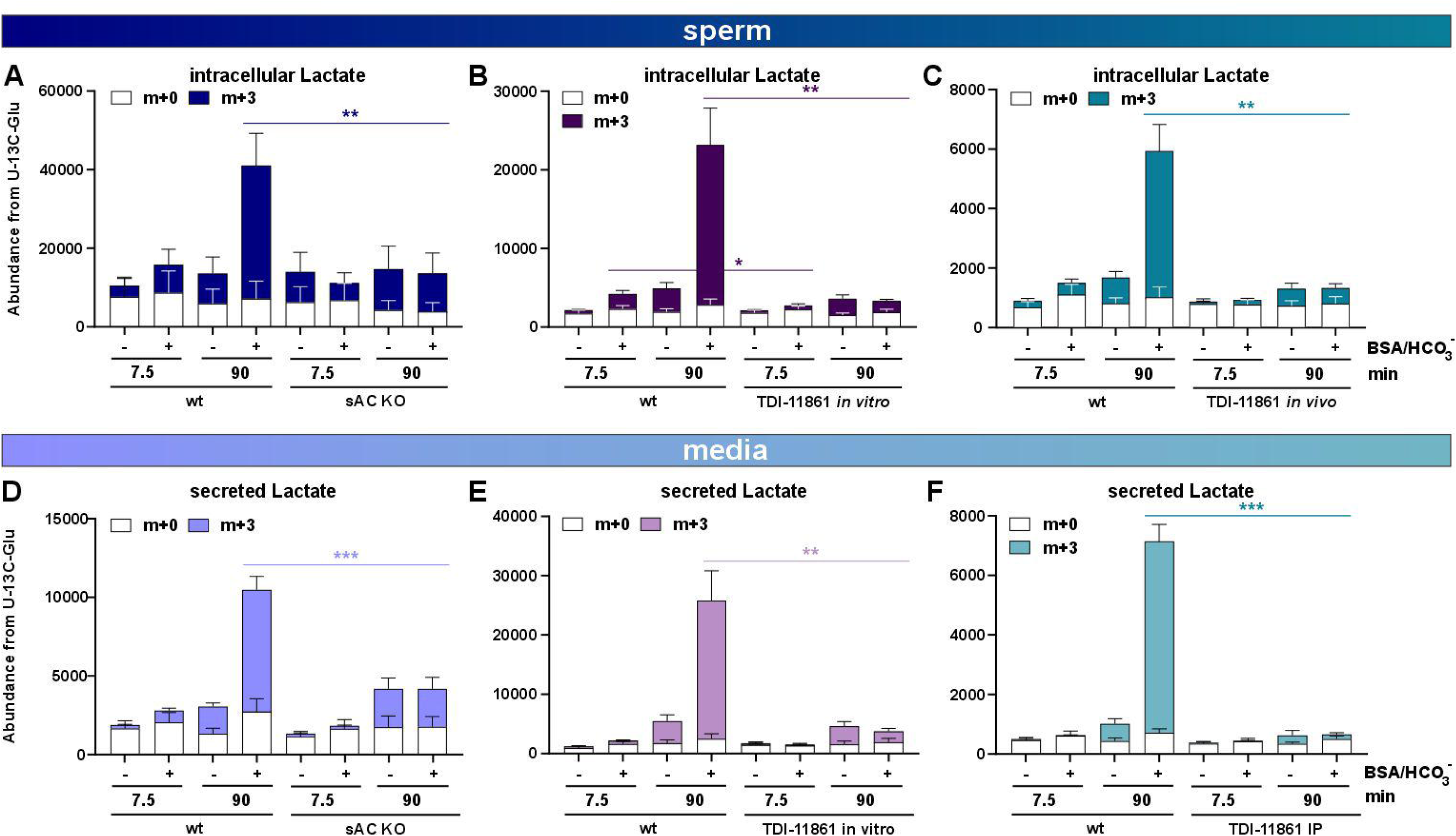
Lactate accumulation and secretion is dependent upon capacitation-induced signaling. **(A-C)** Total ion intensities of unlabeled (^12^C, m+0) and labeled (^13^C, m+3 isotopologues) intracellular lactate in non-capacitating and capacitating **(A)** wild type (wt) and sAC KO sperm, **(B)** control and sAC-inhibited sperm incubated in 100 nM TDI-11861, and **(C)** sperm from vehicle- and TDI-11861-injected mice in U-^13^C-glucose. **(D-F)** Total ion intensities of unlabeled (^12^C, m+0) and labeled (^13^C, m+3 isotopologues) lactate in the media surrounding non-capacitating and capacitating **(D)** wild type (wt) and sAC KO sperm, **(E)** control and sAC-inhibited sperm incubated in 100 nM TDI-11861, and **(F)** sperm from vehicle- and TDI-11861-injected mice in U-^13^C-glucose. Mean + SEM, n=7. Differences between wild-type and capacitation-deficient sperm incubated in non-capacitating or capacitating conditions at the same time point were analyzed using two-tailed, unpaired *t*-test, *P<0.05, **P<0.01, ***P<0.001, ****P<0.0001.

### Sperm do not metabolize extracellular lactate into pyruvate or TCA cycle metabolites

Finding lactate in the media prompted us to examine whether monocarboxylate transporters in the sperm midpiece take up the secreted lactate and metabolize it through the TCA cycle or convert three-carbon metabolites into hexoses. Indeed, U-^13^C-lactate rapidly entered sperm (Fig. S1); however, neither non-capacitating nor capacitating sperm metabolized extracellular lactate into pyruvate, citrate, or other 3- or 6-carbon glycolytic intermediates. Hexose phosphate, PEP, and citrate levels decreased over time confirming the sperm were alive and metabolically active (Fig. S1). As a control, we used somatic HEK293 to confirm that we could observe metabolic utilization of exogenous lactate using our methodology. In HEK293 cells, we observed accumulation of m+3 pyruvate and low levels of m+2 citrate in a time-dependent manner, revealing that in contrast to sperm, HEK293 cells convert extracellular lactate into pyruvate and metabolize it into citrate (Fig. S2).

### Phosphofructokinase is a rate-limiting step of sperm glycolysis

The rate of labeling for the next metabolite after glucose-6-phosphate and fructose-6-phosphate in the glycolytic pathway, fructose-1,6-bisphosphate, was strikingly different from the nearly complete labeling of the hexose phosphates described above. In contrast to the rapid first steps of glycolysis, the accumulation of m+3 and m+6 fructose-1,6,-bisphosphate (F1,6BP) showed a slower kinetic in both capacitating and non-capacitating sperm (Fig. 4A,F). Initially, we only observed a labeling percentage of about 25% F1,6BP in both non-capacitating and capacitating sperm, and a labeling percentage similar to what was observed for hexose phosphate was only achieved at 90 min (Fig. 4A,F; Fig. 1B,C). These data suggest that the activity of phosphofructokinase (PFK), the enzyme converting fructose-6-phosphate into F1,6BP, is rate-limiting glycolytic flux in both capacitating and non-capacitating sperm. We did observe a difference in sperm incubated in capacitating conditions at later time points. After incubating in U-^13^C-glucose for 60 minutes, we detected significantly higher amounts with greater fractional labeling of F1,6BP in capacitating relative to non-capacitating sperm (Fig. 4A,F). Subsequent glycolytic intermediates up to pyruvate also accumulated to greater levels in capacitated sperm at later times (Fig. 4G-J). We presume these increased levels of glycolytic intermediates observed after prolonged times in capacitation media reflect surplus flow through glycolysis as a consequence of the previously described capacitation-induced increase in glucose uptake^6^.

**Fig. 4:**
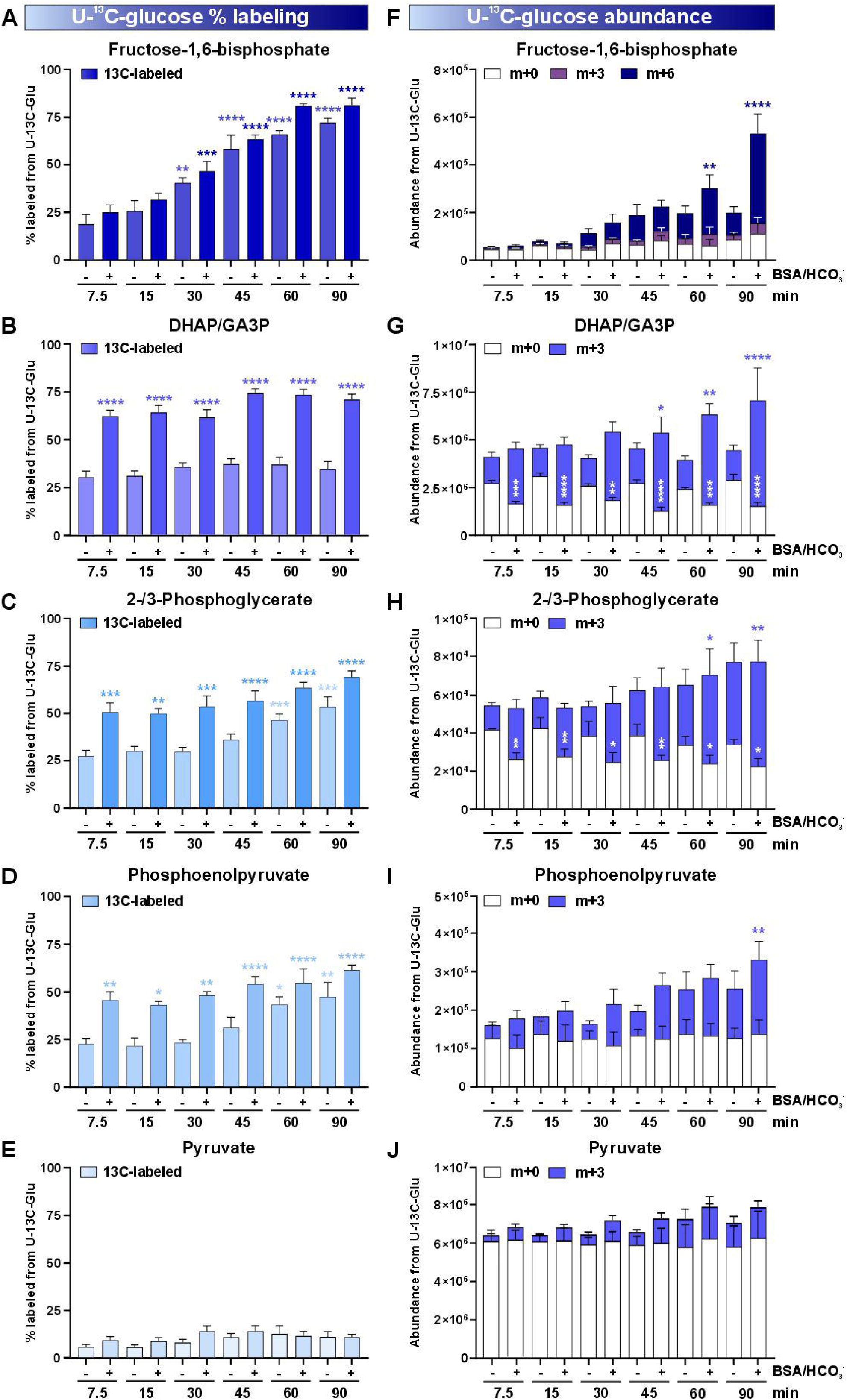
Aldolase regulates glucose flux through glycolysis. **(A-E)** Fractional enrichment of total labeled (^13^C, m+3 or m+6 isotopologues) **(A)** fructose-1,6-bisphosphate, **(B)** DHAP/GA3P, **(C)** 2-,3-phosphoglycerate, **(D)** phosphoenolpyruvate, and **(E)** pyruvate in non-capacitating and capacitating mouse sperm in U-^13^C-glucose. **(F-J)** Total ion intensities of unlabeled (^12^C, m+0) and labeled (^13^C, m+3 or m+6 isotopologues) **(F)** fructose-1,6-bisphosphate, **(G)** DHAP/GA3P, **(H)** 2-,3-phosphoglycerate, **(I)** phosphoenolpyruvate, and **(J)** pyruvate in non-capacitating and capacitating mouse sperm in U-^13^C-glucose. Mean + SEM, n≥4. Differences between conditions were analyzed using one-way ANOVA compared to sperm incubated in non-capacitating conditions at 7.5 min (white asterisk: m+0, blue asterisk: m+3/m+6), *P<0.05, **P< 0.01, ***P<0.001, ****P<0.0001.

### Aldolase is responsible for the capacitation-induced increased glycolytic flux

F1,6BP is cleaved into dihydroxyacetone phosphate (DHAP)/glyceraldehyde-3-phosphate (GA3P) by aldolase. At the earliest times measured, we observed an immediate, capacitation-dependent increase in fractional labeling of DHAP/GA3P (which can’t be distinguished in our chromatographic method) (Fig. 4B). This capacitation-dependent increase was also observed in the subsequent metabolites of lower glycolysis, 2-/3-phosphoglycerate (2-/3-PG), and phosphoenolpyruvate (PEP), which more than doubled in capacitating sperm relative to non-capacitating sperm (Fig 4C,D). These data reveal that an early event during sperm capacitation is an increase in aldolase activity, the enzyme responsible for the conversion of F16BP into DHAP and GA3P. However, we did not detect a capacitation-induced increase in aldolase activity in sperm lysates (Fig. S3). Importantly, performing enzymatic assays in sperm lysates misses allosteric inhibitors and potential changes in aldolase location which could be alternative mechanisms to potentiate aldolase activity.

In sAC KO and sAC-inhibited sperm, the fraction of labeled hexose phosphate and F1,6BP at 7.5 min were comparable to the non-capacitating and capacitating wild type, uninhibited sperm (Fig. 5A,C,E). In contrast, the capacitation-induced increases in labeled DHAP/GA3P and PEP, similar as observed for intracellular and secreted lactate (Fig. 3) were absent in sperm devoid of sAC activity (Fig. 5). The capacitation-induced increases in abundance of labeled F1,6BP, DHAP/GA3P, and PEP at later times were also absent when sAC was knocked out or blocked (Fig. 5B, D, F). These data indicate that capacitating sperm undergo a cAMP-dependent increase in aldolase activity which stimulates flux through glycolysis at the earliest times measured and results in accumulation of glycolytic intermediates at later time points.

**Fig. 5:**
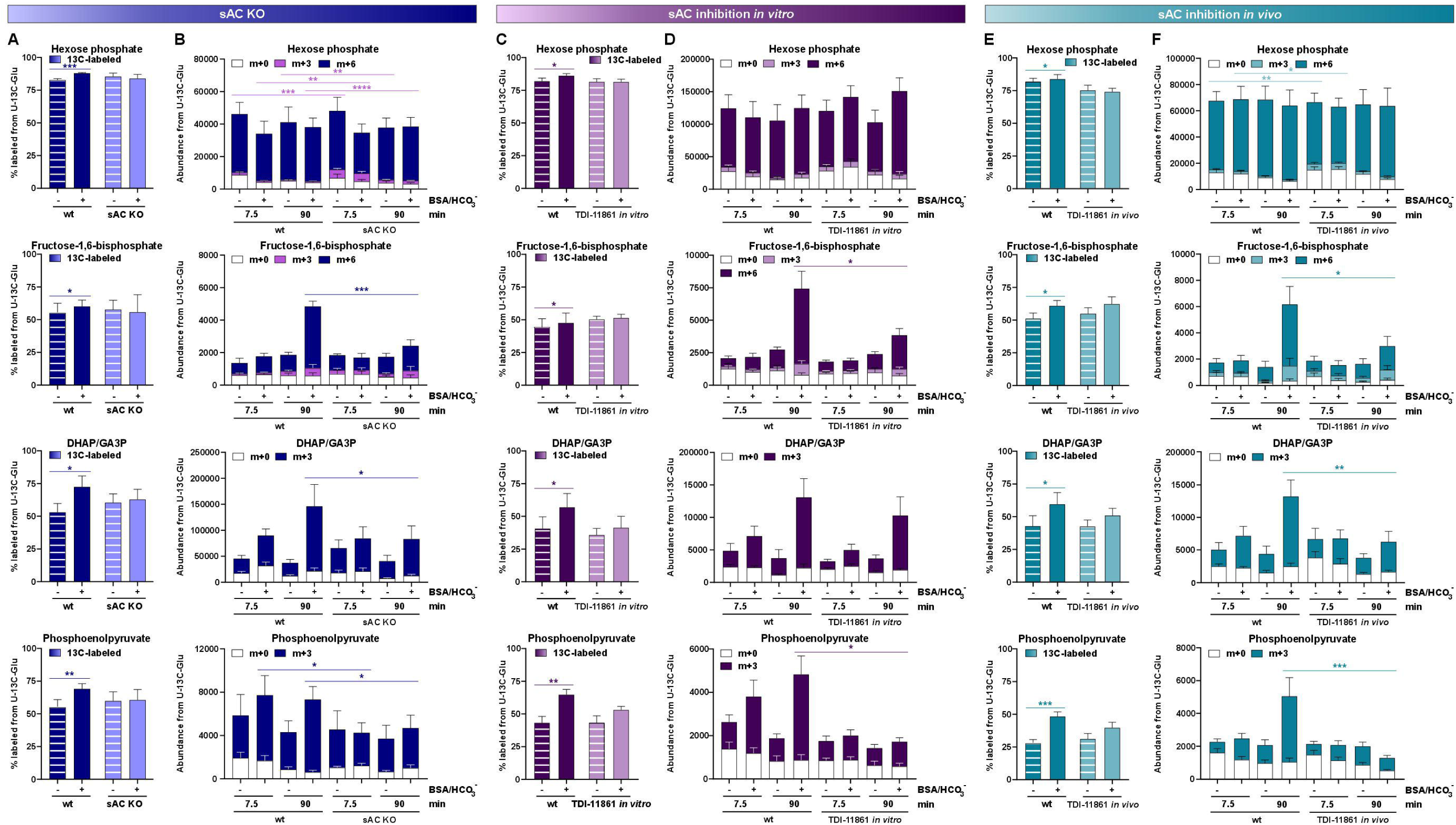
Aldolase regulation of glycolytic flux is dependent upon capacitation-induced signaling. **(A,C,E)** Fractional enrichment of total labeled (^13^C, m+1 to m+6 isotopologues) at 7.5 min and **(B,D,F)** total ion intensities of unlabeled (^12^C, m+0) and labeled (^13^C, m+3 or m+6 isotopologues) glycolytic metabolites in non-capacitating and capacitating **(A,B)** wild type (wt) and sAC KO sperm, **(C,D)** control and sAC-inhibited sperm incubated in 100 nM TDI-11861, and **(E,F)** sperm from vehicle- and TDI-11861-injected mice in U-^13^C-glucose. Mean + SEM, n=4. Differences between non-capacitating and capacitating sperm (A,C,E) and wild-type and capacitation-deficient sperm incubated in non-capacitating or capacitating conditions at the same time point (B,D,F) were analyzed using two-tailed, unpaired *t*-test, *P<0.05, **P<0.01, ***P<0.001, ****P<0.0001.

### Sperm possess two distinct pools of pyruvate

Overall, the fractional labeling of pyruvate was much lower than for the other glycolytic metabolites (Fig. 5I. This disparity can be understood when considering the abundance of labeled and unlabeled pyruvate (5J). Sperm harbor a considerable pool of pyruvate, which remains unlabeled because it is independent from the pyruvate arising from the labeled glucose generated via glycolysis. While the labeled pyruvate generated through glycolysis is rapidly converted into its downstream metabolites preventing the accumulation of m+3 pyruvate, the amount of unlabeled endogenous pyruvate remained constant between non-capacitating and capacitating sperm suggesting it resides in a separate compartment, untouched by the capacitation-induced changes in central carbon flux.

### Sperm metabolize glucose into citrate

In sperm, the purported compartmentalization between glycolysis and oxphos elicits questions about the linkage between the two pathways, specifically whether glycolytically produced pyruvate is converted into citrate. Extracellular flux analysis suggested that the two pathways are connected^7^, and NMR studies confirmed that sperm generate ^13^C-citrate from ^13^C-glucose^34^. SIL with U-^13^C-glucose resulted in m+2 citrate in non-capacitating and capacitating sperm at 7.5 min (Fig. 6B). The conversion of ^13^C-labeled metabolites into citrate was potentiated by capacitation. M+2 citrate was gradually replaced by isotopologues with higher mass, confirming flux through the TCA cycle. We detected not only m+4 and m+6 but also m+3 and m+5 citrate isotopologues. The presence of these various isotopologues of citrate indicate that glycolytically generated pyruvate is converted into m+2 acetyl-CoA by pyruvate dehydrogenase as well as carboxylated into oxaloacetate via pyruvate carboxylase, which then condenses with acetyl-CoA to form citrate (Fig. 6A). In sum, these data confirm that products from exogenous glucose enter the TCA cycle, proving that glycolysis and oxphos are linked in sperm and that there is increased flux through glycolysis and oxphos in capacitating sperm. While our data don’t exclude that pyruvate generated by glycolysis travels from the principal piece to the midpiece, the rapid metabolism of ^13^C-pyruvate and lack of metabolism of exogenous ^13^C-lactate rather suggests that glycolysis is also happening in the midpiece in close proximity to the mitochondria. Interestingly, we also detected significantly increased amounts of unlabeled citrate when sperm were subjected to capacitating conditions (Fig. 6B); therefore, capacitation also potentiates the metabolism of internal carbon sources into the TCA cycle.

**Fig. 6:**
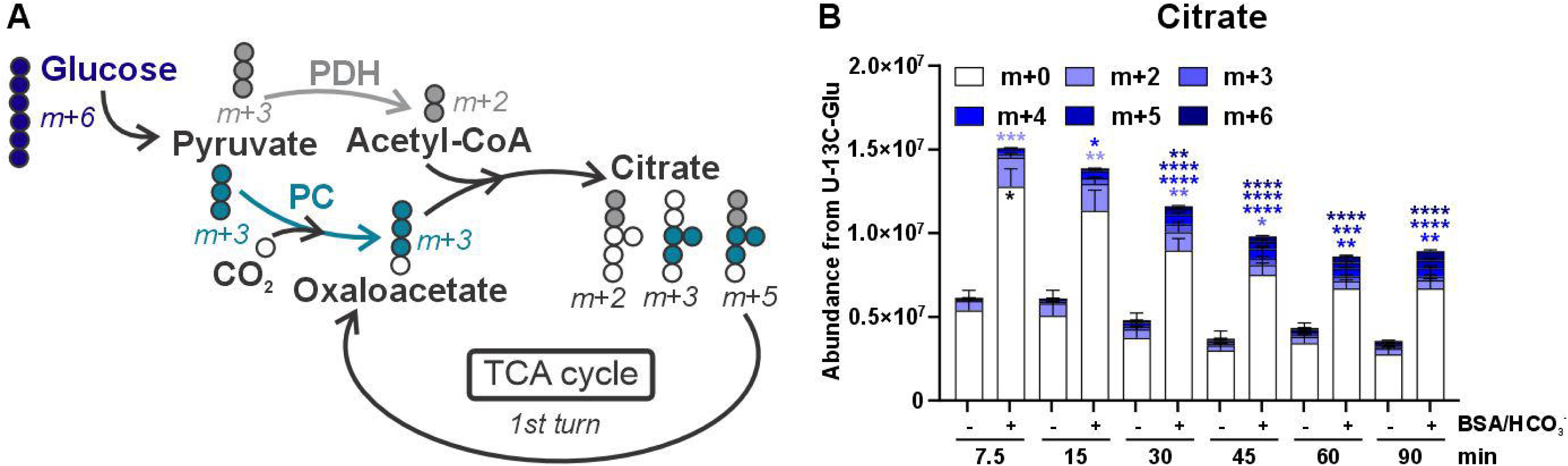
Glucose metabolism results in rapid citrate labeling. **(A)** Schematics of metabolic flux into citrate from U-^13^C-pyruvate derived from U-^13^C-glucose. Pyruvate enters the TCA cycle either through conversion into acetyl-CoA by pyruvate dehydrogenase (PDH) or through conversion into oxaloacetate by pyruvate carboxylase (PC) initially resulting in m+2, m+3, and m+5 citrate isotopologues. **(B)** Total ion intensities of unlabeled (^12^C, m+0) and labeled (^13^C, m+2 to m+6 isotopologues) citrate in non-capacitating and capacitating sperm incubated for the indicated time points in 5.6 mM U-^13^C-glucose. Mean + SEM, n=8. Differences between conditions were analyzed using one-way ANOVA compared to sperm incubated in non-capacitating conditions at 7.5 min (blue asterisk: m+2-m+6), *P<0.05, **P< 0.01, ***P<0.001, ****P<0.0001.

### Sperm capacitation reduces flux through the pentose phosphate pathway

We next examined the various intermediates leading from hexose phosphates through glycolysis to ask which step is regulated in capacitating sperm. The first branch point in glucose metabolism is the fate of G6P; i.e., whether it is isomerized into F6P to proceed through glycolysis or oxidized into 6-phosphogluconolactone via glucose-6-phosphate dehydrogenase (G6PD) to enter the pentose phosphate pathway (PPP). G6PD is present in mouse sperm^24^, and we observed m+5 pentose phosphate metabolites (phosphorylated pentoses are not distinguished in our chromatographic method and are therefore collectively referred to as pentose phosphate) at the first time points analyzed in both non-capacitating and capacitating sperm confirming that the PPP is active in mouse sperm (Fig. 7). Surprisingly, while the unlabeled pool of pentose phosphates remained unchanged over time in both capacitating and non-capacitating sperm, the fractional labeling and total amount of m+5 pentose phosphates were higher in non-capacitating than in capacitating sperm (Fig. 7A,B). These data suggest that resting epididymal mouse sperm flux carbon through the PPP at higher levels than capacitating sperm. Stored epididymal sperm may need to flux carbon through the PPP to generate more anti-oxidative reducing equivalents while they are dormant in the epididymis awaiting ejaculation. In contrast, when capacitation is induced, carbon flux through the PPP is reduced either passively due to increased glycolytic activity or actively by decreasing G6PD activity. Reducing the flux through the PPP allows capacitating sperm to shunt more exogenous carbohydrates through glycolysis for increased ATP generation.

**Fig. 7:**
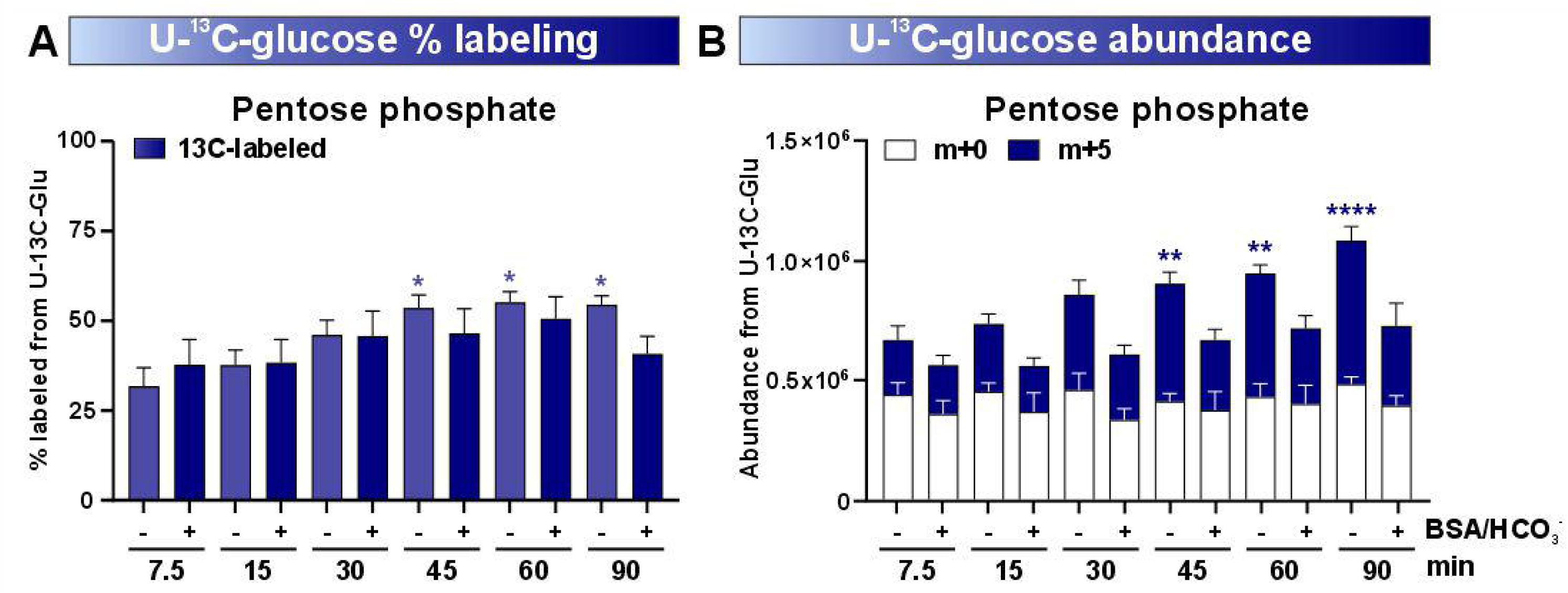
Capacitation-induced increase in glycolytic flux reduces flux through the pentose phosphate pathway. **(A,B) (A)** Fractional enrichment of total labeled (^13^C, m+5 isotopologue) and **(B)** total ion intensities of unlabeled (^12^C, m+0) and labeled (^13^C, m+5 isotopologues) pentose phosphate in non-capacitating and capacitating epididymal mouse sperm incubated for the indicated time points in 5.6 mM U-^13^C-glucose. Mean + SEM, n≥4. Differences between conditions were analyzed using one-way ANOVA compared to sperm incubated in non-capacitating conditions at 7.5 min; *P<0.05, **P< 0.01, ****P<0.0001.

## Discussion

Stable isotope labeling is a key approach for understanding the dynamics of metabolic reprogramming in mammalian cells. SIL in sperm is challenging due to their small cytoplasmic volume and heterogeneous sample composition resulting in low metabolite abundances. Previous studies employing SIL to investigate mammalian sperm metabolism^38–43,44^ included a limited repertoire of metabolites and focused exclusively on non-capacitated sperm^38–43,44^. Our studies compared cauda epididymal mouse sperm, which resemble the metabolic state of dormant stored sperm, with sperm capacitated *in vitro*. We also combined highly concentrated sperm samples with mM concentrations of ^13^C-carbohydrates, ion-pair chromatography, and sensitive mass spectrometry instrumentation which allowed us to follow the distribution of labeled carbon atoms into most glycolytic metabolites, as well as pentose phosphates and citrate. Thus, our studies extend the exploration of mammalian sperm metabolism to the dynamic changes of the full central carbon metabolic network during capacitation-induced metabolic reprogramming. We confirmed the increase in glycolytic flux and conversion of glycolysis-generate pyruvate into citrate and secreted lactate during mouse sperm capacitation and dissected the rate changes of individual glycolytic enzymes. Importantly, we identified aldolase as the enzyme responsible for controlling the capacitation-induced increase in glycolytic flux.

Similar to many tissues and cell types, PFK is a rate-limiting step of glycolytic flux in sperm, but the observed time-dependent increase in PFK activity is not responsible for the capacitation-induced stimulation of glycolytic flux. Instead, we find that sperm utilize a second enzymatic step, aldolase, to respond to the increased energy demand of motility through the female reproductive tract and capacitation-associated processes. Little information is available if aldolase plays a similar role for glycolytic flux regulation in somatic cells. Overexpression of ALDOA in pancreatic and other cancer types increases glycolytic rate, which correlates with poor prognosis and survival rates (reviewed in^45^). However, stimulating aldolase activity via increased expression is not possible in sperm because epididymal sperm are transcriptionally and translationally silent. Mice express three sperm specific isoforms of ALDOA with unique N-terminal extensions that are predicted to be responsible for tight binding to the fibrous sheath^46,47^. The aldolase-mediated increase in DHAP/GA3P accumulation was blocked when capacitation was prevented by knockout or inhibition of sAC revealing that capacitation-induced signal transduction is responsible for stimulating aldolase activity. Among the molecular changes associated with sperm capacitation is a sAC-dependent increase in tyrosine phosphorylation^14^, and 2D gel electrophoresis combined with mass spectrometry identified testis-specific aldolase as a substrate for tyrosine phosphorylation^48^. However, we and others^48^ also showed that the *in vitro* activity of aldolase was unchanged in capacitating sperm relative to non-capacitated sperm. Therefore, capacitation-dependent changes are not reflected by changes to the enzymatic aldolase activity in sperm extracts, and we hypothesize that capacitation causes stimulation of aldolase activity due to a change in localization, conformation, or accessibility to its substrate F1,6BP.

The unique compartmentalization of metabolic machinery into distinct sections of the sperm tail (i.e., glycolytic enzymes predominantly in the principal piece with mitochondria and oxphos machinery exclusively in the midpiece) fueled suspicion that sperm do not feed glycolytic products directly into the mitochondria for oxphos. Data presented here, along with published studies demonstrating that capacitation stimulates glucose uptake into the midpiece and capacitation-dependent increases in oxphos are dependent upon glycolysis^7^, reveal that the midpiece also contains glycolytic machinery to metabolize the locally imported glucose into pyruvate which feeds mitochondrial oxphos. The pyruvate generated from exogenously supplied glucose does not accumulate or appear to persist for great lengths of time suggesting it is utilized rapidly. While our previous studies^34^ showed that exogenously supplied glucose is metabolized into citrate, here we extend these results to show that exogenous glucose generates each of the isotopologues of citrate confirming it is cycling through the TCA cycle. The rapid metabolism of glycolysis-generated pyruvate and absence of metabolism of exogenous lactate points towards glycolytic activity in the midpiece in close proximity to the mitochondria.

If entry into the mitochondrial TCA cycle and oxphos is the consequence of glycolytically generated pyruvate in the midpiece, we assume the high levels of lactate accumulating and secreted from sperm are generated in the principal piece, where glycolytic machinery is abundant and there are no mitochondria. Germ cells express a specific isoform of lactate dehydrogenase, LDHC, whose absence leads to male infertility along with decreased ATP levels and sperm motility^49–51^. LDHC favors reducing pyruvate generating lactate while also regenerating NAD^+^ over the reverse reaction to oxidize lactate^52^, and as proposed by Storey and Visconti^27,32^, regenerating NAD^+^ ensures that glycolytic enzymes are not NAD^+^-limited while yielding a product (lactate) that can easily be secreted.

Analyzing extracellular metabolites in sperm media revealed that secretion of lactate is a consequence of sperm glycolysis. Since we did not detect ^13^C-pyruvate when incubating sperm with ^13^C-lactate our data suggest that secreted lactate is not metabolically utilized by mouse sperm. Schmidt *et al.*, however, report the highest level of mitochondrial respiration with extracellular lactate^48^. Additional experiments are required to explain the discrepancy between our studies. The elevated production and subsequent secretion of lactate may suggest that lactate is simply a waste product of carbon metabolism, which would not be surprising considering sperm do not utilize carbons for generating biomass. In this case, lactate secretion from sperm together with H^+^, would prevent intracellular acidification to ensure continued high rates of glycolysis^49,50^. It remains possible that secretion of lactate might have functional relevance for sperm as they transit the female reproductive tract. For example, secreted lactate may fulfill a role as immunosuppressor as proposed for tumor cells^51^.

While glucose-generated pyruvate is rapidly converted into acetyl-CoA or oxaloacetate resulting in labeled citrate in the midpiece and into lactate in the principal piece, the majority of pyruvate found in sperm is stored pyruvate, not generated from exogenously supplied glucose. It is tempting to hypothesize that this pool of pyruvate may exist in the midpiece, where it could directly feed mitochondrial oxphos as an internal, stored pool of energy. In fact, in capacitating sperm, the majority of generated citrate is unlabeled, indicating that it derives from an internal, stored energy source. The greatest level of capacitation-dependent citrate produced is seen at the earliest time measured (7.5 minutes), which means that the major consumption of stored pyruvate may already have occurred by this time, which might explain why the stored pyruvate does not appear to diminish at later time points. However, it is also possible that sperm generate citrate from other internal energy stores not measured in our studies (i.e., glutamine or free fatty acids). Free fatty acids are a particularly attractive potential source of stored energy in sperm. Supporting this hypothesis, defective fatty acid oxidation negatively affects sperm function^44^.

Utilizing SIL with U-^13^C-glucose to its full capacity combined with genetic and pharmacological tools allowed us to provide insights into mammalian sperm metabolic reprogramming induced upon ejaculation. These insights support a model for the metabolic shifts which occur in sperm which are tailored to ensure their ability to reach and fertilize the oocyte (Fig. 8). Prior to ejaculation, sperm are stored in the cauda epididymis in a dormant state. These resting sperm utilize glucose to produce energy via glycolysis and oxphos, but they also shunt significant amounts of glucose into the pentose phosphate pathway to produce reducing equivalents to ensure protection of cellular machinery at the expense of ATP generation. Once sperm are ejaculated, they begin swimming and initiate capacitation. To meet the increased energy demands of these essential processes, sperm increase glucose uptake and stimulate aldolase activity to increase flux through glycolysis. The pentose phosphate pathway remains active; however, flux through the PPP is reduced in capacitating sperm in favor of flux through glycolysis. In the principal piece, where glycolysis represents the main energy producing metabolic process supporting motility, pyruvate produced from glycolysis is reduced to lactate, which is secreted to prevent intracellular acidification while simultaneously regenerating NAD^+^ to support upstream glycolytic reactions. In contrast, in the midpiece, glycolytically generated pyruvate enters mitochondria where it feeds the TCA cycle and oxphos, which are more efficient at producing ATP. Among the changes during capacitation, sperm alter their motility pattern to hyperactivated motility. Others demonstrated that hyperactivated motility is dependent upon mitochondrial oxphos^53^, and in addition to being supported by exogenous glucose, via glycolysis, mitochondrial oxphos is also fed by endogenous energy sources, which may be stored pyruvate or possibly free fatty acids. Capacitation also stimulates the utilization of these endogenous energy sources.

**Fig. 8:**
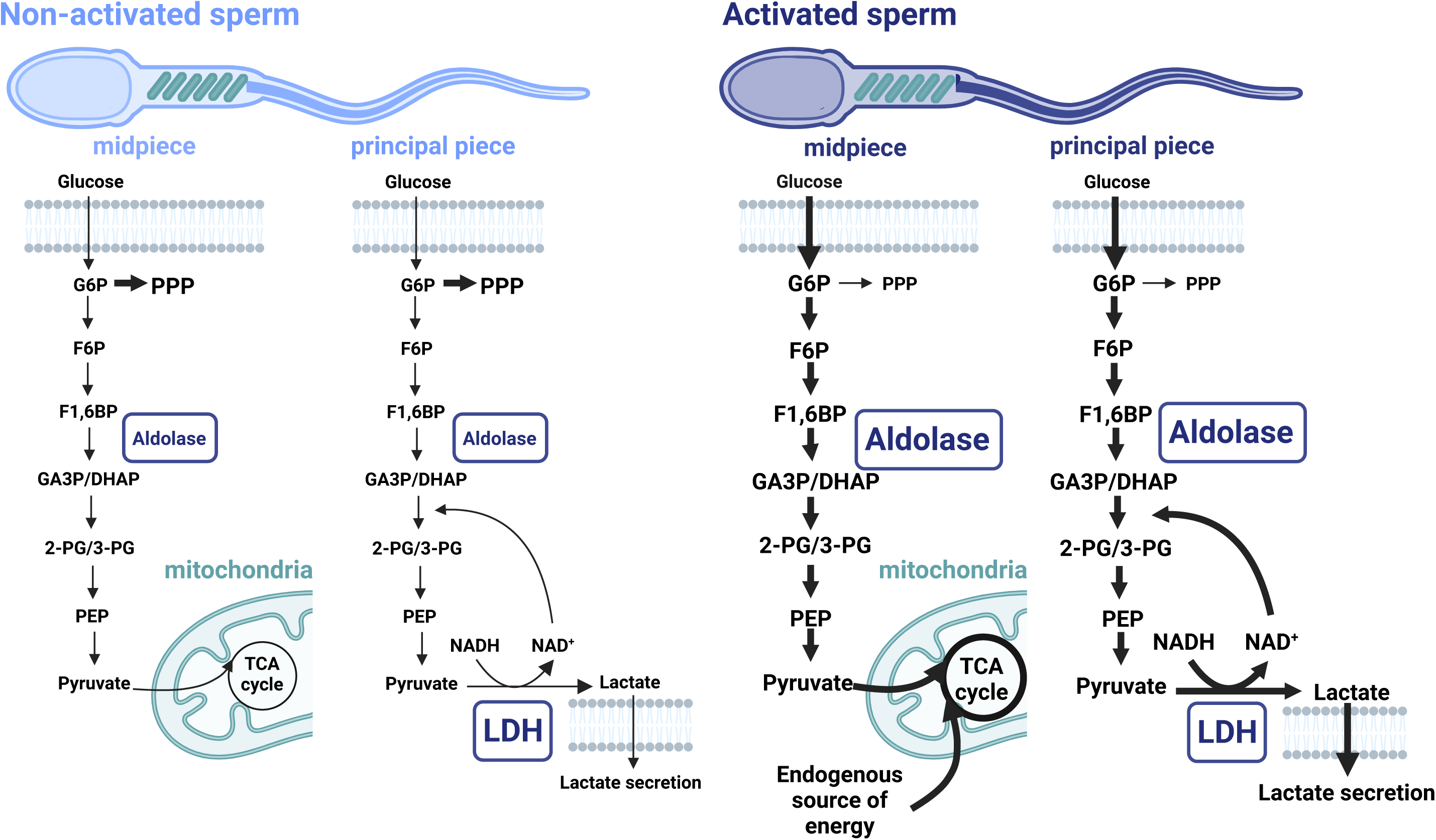
Model of capacitation-induced changes in sperm central carbon metabolism. In capacitating sperm, glucose flux through glycolysis is increased at the expense of the pentose phosphate pathway. Increased flux through glycolysis is due to activation-induced stimulation of aldolase activity. In the mitochondria-containing midpiece, glycolytically generated pyruvate feeds the TCA cycle. In the mitochondria-free principal piece of the tail, pyruvate produced from glycolysis is reduced to lactate and secreted. A yet unknown endogenous source of energy is also feeding the upregulation of TCA cycle intermediates.

## METHODS

### Reagents

Reagents for media preparation were purchased from Sigma-Aldrich. DMSO, PEG 400, BSA, HCO_3_ ^−^, MeOH, MtBE, TBA, mPIC, and PhosStop were purchased from Sigma-Aldrich, ^13^C-labeled tracer from Cambridge Isotope Laboratories, and PBS, DMEM, and FBS from Corning. TDI-11861 was dissolved in DMSO resulting in a final concentration of 0.1 % DMSO per sample. In every experiment, 0.1 % DMSO was included as vehicle control.

### Mice

Adult 8-10 week old C57BL/6N (Stock #: 027) mice were purchased from Jackson Laboratory and allowed to acclimatize before use. *Adcy10* KO^22^ and their corresponding wild type were in the C57BL/6J background and bred in-house. Animal experiments were approved by Michigan State University’s and Weill Cornell Medicine’s Institutional Animal Care and Use Committees (IACUC).

### Sperm preparation

Epididymal mouse sperm were isolated by incision of the cauda epididymis in 500 µl modified TYH medium (in mM: 135 NaCl, 4.7 KCl, 1.7 CaCl2, 1.2 KH2PO4, 1.2 MgCl2, 10 HEPES, pH 7.4 adjusted at 37°C with NaOH) with the respective energy source, prewarmed at 37°C. After 15 min swim-out at 37°C, sperm were counted using a hematocytometer and sperm cell numbers were adjusted to a concentration of 1×10^7^ cells/ml. Sperm were washed twice with the TYH buffer with the respective energy source used for the experiment (glucose, or lactate (0.56 mM) by centrifugation at 700xg for 5 min. For capacitation, sperm were incubated in TYH containing 3 mg/ml BSA and 25 mM NaHCO ^−^ in a 37°C incubator.

### Sperm ^13^C stable isotope labeling and extraction

^13^C metabolites were analyzed in the MSKCC, MSU, and Van Andel metabolomics cores to make use of different standard libraries and to confirm results. Specifics for metabolite extraction and analysis at MSKCC and Van Andel are included as supplemental information S4. Fractional enrichment results for overlapping metabolites were consistent between all facilities. Graphs showing fractional enrichment show averages from data generated in all facilities, plots showing relative abundance summarize data generated at MSU.

Epididymal sperm from wild type or sAC KO mice were isolated in non-capacitating TYH buffer with 0.56 mM glucose or lactate, pooled and counted. Aliquots of 50 ul sperm were added to 450 μl non-capacitating or capacitating TYH containing 0.56 mM glucose or lactose to a final concentration of 5 x 10^6^ sperm/ml. 5 μl of 560 mM U-^13^C-glucose or U-^13^C-lactate or the corresponding ^12^C control were added. To block sperm capacitation *in vitro*, non-capacitating and capacitating sperm were incubated in the presence of 100 nM TDI-11861 for the duration of SIL. For SIL after inhibition of capacitation *in vivo*, 150 µl of solution containing 50 mg/kg TDI-11861 were administered systemically via intraperitoneal injection; control males were subjected to 150 µl vehicle control (DMSO:PEG 400:PBS 1:4:5 (v/v) as described in^5^. Sperm were isolated one hour later and SIL was performed as described above.

At the indicated time points, sperm and supernatant were separated by centrifugation. 450 μl of supernatant were transferred to a new vial and 100 μl of the supernatant were mixed with 400 % ice-cold MeOH. Media without sperm was used as control and processed under the same conditions.

Sperm pellets were washed with 500 μl cold 0.9 % NaCl, and 500 μl ice-cold MtBE:MeOH (3:1) was added. Samples were immediately vortexed and placed on dry ice. Sperm were sonicated for 10 min in a water-bath sonicator, mixed with 300 μl H2O:MeOH (3:1) and vortexed again. Samples were centrifuged at 16,000xg for 10 min at 4°C and 450 μl of the lower polar phase (MSU) or MeOH (MSKCC) was transferred to new tubes. Sperm and supernatant samples were centrifuged at 20,000xg for 20 min at 4°C and 450 μl of supernatant were transferred to a fresh tube, dried in a vacuum evaporator and stored at −80°C until analysis. Before LC-MS/MS analysis, samples were resuspended in 30 μl mobile phase A and incubated on wet ice for 20 min while being vortexed every 5 min. Samples were centrifuged at 20,000 x g for 20 min at 4°C and 25 μl of the supernatant was transferred into LC vials for injection.

### HEK293 cell ^13^C stable isotope labeling and extraction

Human kidney epithelial HEK293 cells (ATCC CRL-1573), were maintained in Dulbecco’s modified eagle medium (DMEM) high glucose media with glutamine, pyruvate, and phenol red (Gibco) and 10% fetal bovine serum. Cells were maintained at 37 °C in a 5% CO2 incubator. For metabolite extraction, cells were seeded at a density of 2 × 10^6^ onto 10 cm plates. At a confluency of 70-80 %, cells were washed with PBS and 5 ml DMEM without energy substrate, and phenol red, substituted with 0.56 mM lactate was added. 50 μl of 560 mM U-^13^C-lactate or the corresponding ^12^C control was added. At the indicated time points, 300 ul of the supernatant was mixed with 1.2 ml ice-cold 100 % MeOH. Media without cells was used as control and processed under the same conditions. The rest of the media was removed, cells were washed with 0.9 % ice-cold NaCl, and 3 ml ice-cold 80% MeOH was added. Plates were stored in a −80 °C freezer for at least 30 min. Cells were scraped off the plate and cells and the organic solvent was collected in microcentrifuge tubes. Samples were sonicated for 10 min in a water-bath sonicator, centrifuged at 1600xg for 10 min at 4°C, and 1.4 ml of organic solvent was transferred to new tubes. Sperm, supernatant, and media samples were dried in a vacuum evaporator and stored at −80°C until analysis. Before LC-MS/MS analysis, samples were resuspended in 60 μl mobile phase A and incubated on wet ice for 20 min while being vortexed every 5 min. Samples were centrifuged at 20,000xg for 20 min at 4°C and 50 μl of the supernatant was transferred into LC vials for injection.

### LC-MS analysis of ^13^C metabolites

Metabolites were analyzed for relative abundance by high-resolution accurate mass detection using a Xevo G2-XS Qtof mass spectrometer (Waters) coupled to a Waters liquid chromatography systems. An Acquity HSS T3 (1.8 μm, 2.1 mm x 150 mm) column (Waters, Eschborn, Germany) was used for chromatographic separation and the elution gradient was carried out with a binary solvent system. Mobile phase A consisted of LC/MS grade H2O with 3% LC/MS grade methanol, 10 mM tributylamine, and 15 mM acetic (pH 5.0 ± 0.05) and solvent B was LC/MS grade 100% methanol. A constant flow rate of 200 μl/min was maintained and the linear gradient employed was as follows: 0–2.5 min 100% A, 2.5–5 min increase from 0 to 20% B, 5–7.5 min maintain 80% A and 20% B, 7.5–13 min increase from 20 to 55% B, 13–15.5 min increase from 55 to 95% B, 15.5–18.5 min maintain 5% A and 95% B, 18.5–19 min decrease from 95 – 0% B, followed by 6 min of re-equilibration at 100% A. The desolvation temperature was set to 350°C, capillary voltage was set to 2 kV, the sample cone voltage was 35 V, the cone gas flow was 40 l/hr, desolvation gas flow was 800 l/hr, and the source temperature was 100°C. The column temperature was maintained at 25°C and sample volumes of 10 μL were injected. Data were acquired using two acqusition functions. Function 1 was a full-scan method used to acquire data with *m/z* scan range from 50 to 1000 and scan time of 0.4 s. Function 2 employed target enhancement set at mass 87 and data were acquired with m/z scan range from 50 to 200 with a scan time of 0.3 s. Lockmass correction was applied using leucine enkephalin as the reference standard. Instrument control and acquisition was carried out by MassLynx software (Waters).

### Targeted metabolomics data analysis

Peak picking and integration was conducted in MassLynx using in-house curated compound data bases of accurate mass MS1 and retention time derived from analytical standards. Full scan raw data files for all samples of a given experiment were imported and metabolite peaks were auto-integrated based off method-specific, in-house curated compound databases that included molecular formula, precursor adducts, and explicit retention times collected from on-method analyzed chemical standards. Manual peak integrations were performed as necessary to account for any tailing peaks and minor retention time shifting. Natural abundance correction was performed using the R package IsocorrectoR.

### Aldolase Assay

Aliquots of 5×10^6^ sperm were incubated for 90 min in non-capacitating or capacitating TYH buffer with glucose. The sperm pellet was incubated for 10 min on ice in 400 µl lysis buffer (50 mM Tris pH 8.0, 1 mM EDTA-NaOH pH 8.0, 0.1% Triton, mPIC, PhosSTOP, pH 7.4). Enzymatic activity was measured in the supernatant separated from the sperm pellet by centrifugation at 10,000xg for 5 min. To determine aldolase activity a commercial aldolase assay kit (Sigma Aldrich MAK223) was used according to the manufacturer’s instructions.

### Statistical analysis

Statistical analyses were performed using GraphPad Prism 5 (Graph-Pad Software). All data are shown as the mean + SEM. Statistical significance between two groups was determined using two-tailed, paired *t*-tests with Welch’s correction, statistical significance between multiple groups was determined using one-way ANOVA with Dunnett‘s correction after confirming normal distribution via D’Agostino-Pearson and homoscedasticity (Breusch–Pagan test). Differences were considered to be significant if ∗P < 0.05, ∗∗P < 0.01, ∗∗∗P < 0.001, and ∗∗∗∗P < 0.0001.

## DATA SHARING PLAN

### Lead contact

Additional information and requests for reagents and resources should be directed to and will be fulfilled by the lead contact, Melanie Balbach.

### Materials availability

This study did not generate new unique reagents.

### Data and code availability

Metabolomics data are deposited at the Metabolomics Workbench^78^ and will be publicly available as soon as the manuscript is published. Accession numbers will be listed in the key resources table. This paper does not report original code. Any additional information required to reanalyze the data reported in this paper is available from the lead contact upon request.

## ACKNOWLEDGMENT

The authors wish to thank Daniel Vocelle and Ola Skirycz for a critical revision of the manuscript. We are grateful to the Lunt lab for sharing their tissue culture hood and reagents with us and to all of the MSU Mass Spectrometry and Metabolomics Core staff for their support. This study was supported by the Eunice Kennedy Shriver National Institute of Child Health and Human Development grants HD088571 (to J.B., L.R.L., and P.E.V.) and HD-038082 (to P.E.V). Dedicated to my friend Claudia.

## AUTHOR CONTRIBUTIONS

M.B. conceived the project, designed and performed the experiments, analyzed and interpreted data, generated the figures, and wrote the manuscript. S.V. designed and performed the experiments and analyzed and interpreted data. A.K., R.D.S., T.S., L.K., K.P., L.A., M.J., and A.J. performed experiments and analyzed data. L.R.L., J.B., P.E.V., S.V., and J.C. conceived the project, interpreted data, and revised the manuscript critically.

## SUPPLEMENTAL INFORMATION

Document S1. Figure S1-S3, Method details MSKCC and Van Andel

