## Supplemental data for "Sperm meet the elevated energy demands to attain fertilization competence by increasing flux through aldolase"

**Supplementary figures**

**Figure S1**

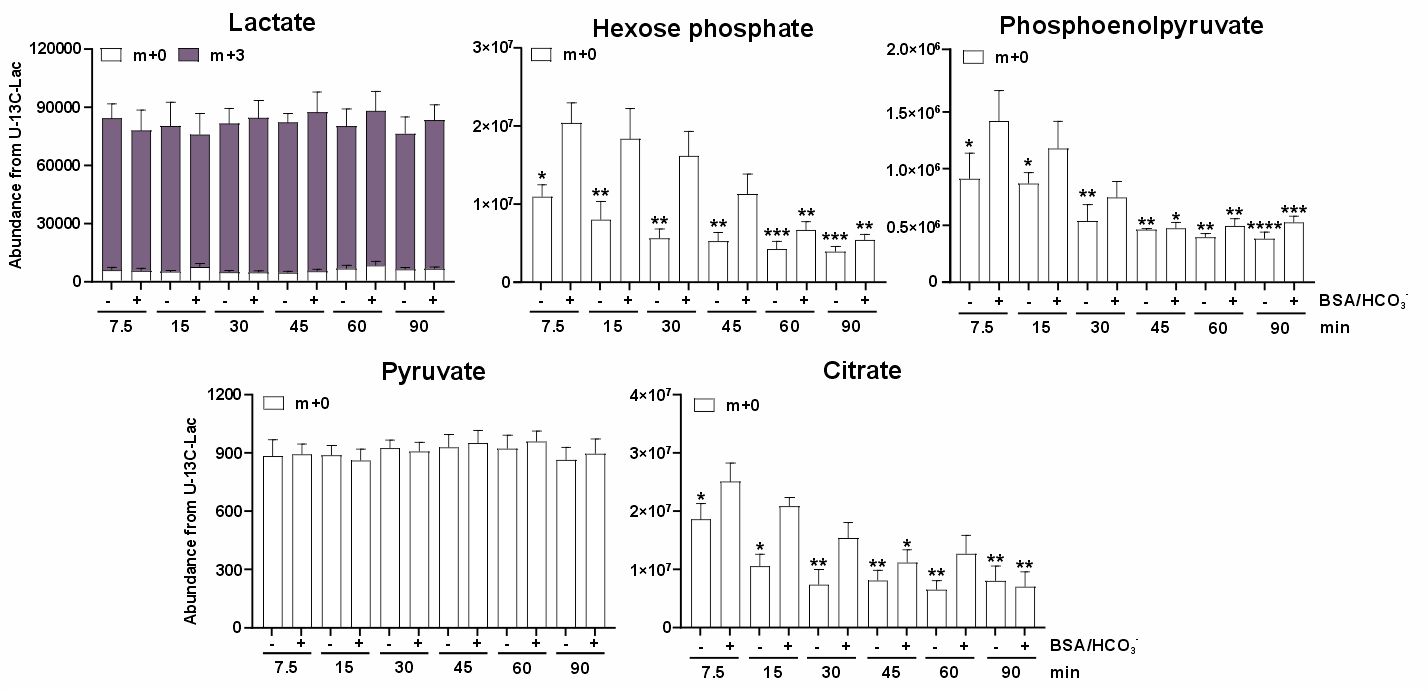

**Figure S1: Sperm take up extracellular lactate but do not metabolize it into glycolytic or TCA cycle intermediates**

Total ion intensities of unlabeled (^12^C, m+0) and labeled (^13^C, m+3 isotopologues) glycolytic metabolites and citrate in non-capacitating and capacitating sperm incubated for the indicated time points in 5.6 mM U-^13^C-lactate. Mean + SEM, n=4. Differences between conditions were analyzed using one-way ANOVA compared to sperm incubated in non-capacitating conditions at 7.5 min, *P<0.05, **P< 0.01, ***P<0.001, ****P<0.0001.

**Figure S2**

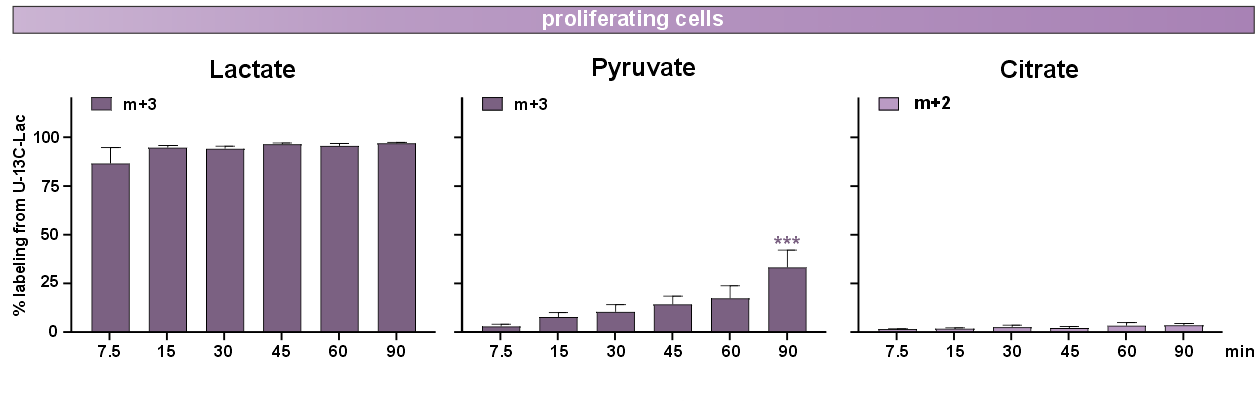

**Figure S2: Somatic HEK293 cells metabolize exogenous lactate**

Fractional enrichment of labeled (^13^C, m+3 isotopologues) lactate, pyruvate, and citrate in HEK293 cells incubated for the indicated time points in 5.6 mM U-^13^C-lactate. Mean + SEM, n=5. Differences between conditions were analyzed using one-way ANOVA compared to HEK293 cells at 7.5 min, ***P<0.001.

**Figure S3**

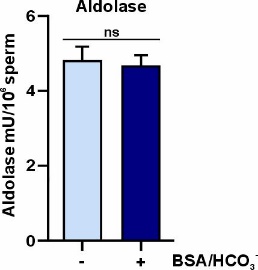

**Fig. S3: Aldolase activity does not increase during sperm capacitation**

Enzymatic activity of Aldolase in non-capacitated and capacitated mouse sperm in 5.6 mM glucose. Differences between non-capacitated and capacitated sperm were analyzed using two-tailed, unpaired *t*-test, ns = not significant.

**Sperm ^13^C stable isotope labeling and extraction (MSKCC and Van Andel metabolomics core)**

At MSKCC, metabolites from sperm pellets were extracted with 450 μl ice-cold MeOH, immediately vortexed and placed on dry ice.

To send sperm pellets to Van Andel, sperm pellets were washed with 500 ul cold 0.9 % NaCl, snap-frozen in liquid nitrogen and shipped on dry ice. At Van Andel, metabolites were extracted from sperm cells using a modified Bligh-Dyer^54^. Briefly, 690 µL of ice-cold 1:1 chloroform:methanol (v/v) was added to pellets of 5x10^6^ cells, vortexed, sonicated for 5 minutes in a water-\bath sonicator, and incubated on ice for 30 min. Then, 310 µl of water was added, vortexed, incubated on ice for 10 min, and centrifuged at 14,000xg for 10 minutes to induce phase separation. 524 µl of the upper, aqueous phase was dried in a vacuum evaporator and resuspended in 50 µl mobile phase A for analysis.

**LC-MS analysis of ^13^C metabolites (MSKCC and Van Andel metabolomics core)**

At MSKCC, metabolites were analyzed by ion pair LC-MS analysis on a 6230 TOF mass spectrometer with dual JetStream source (Agilent Technologies) in negative ionization mode using a Waters XSelect HSS T3 column (150 x 2.1 mm, 3.5 µm particle size), and applying a gradient of solvent A (5 mM octylamine and 5 mM acetic acid in water) and solvent B (5 mM octylamine and 5 mM acetic acid in 90:10 methanol:water), with a post-column flow consisting of a blend of acetone:DMSO (90:10) at 300 µl/min. The analytical gradient was 0-3.5 min, 1% B; 4-15 min, 35% B; 20-22 min, 100% B; 22-27 min, 1% B. Other LC parameters were as follows: flow rate 300 µl/min, column temperature 40°C, and injection volume 5 µl. MS parameters were as follows: gas temp: 250°C; gas flow: 9 L/min; nebulizer pressure: 35 psig; sheath gas temp: 250°C; sheath gas flow: 12 L/min; VCap: 3500 V; and fragmentor: 125 V. Data were acquired from 50 to 1700 m/z with active reference mass correction (m/z: 119.0363 and 966.0007) infused through a second nebulizer according to the manufacturer’s instructions.

At Van Andel Insitute’s Mass Spectrometry Core (RRID:SCR 024903), as previously reported^55-59^, metabolomics was conducted using tributylamine ion-paired liquid chromatography on an Orbitrap Exploris 240. For the chromatography, mobile phase A was LC/MS H_2_O with 3% LC/MS grade MeOH, mobile phase B was LC/MS grade methanol, and both mobile phases contained 10 mM tributylamine, 15 mM acetic acid, and 0.01% medronic acid. For the re-equilibration gradient, mobile phase A was kept the same, and mobile phase B was 99% LC/MS grade acetonitrile. Column temperature was kept at 35ºC, flow rate 0.25 mL/min, and the solvent gradient was as follows: 0-2.5 min held at 0% B, 2.5-7.5 min from 0% B to 20% B, 7.5-13 min from 20% B to 45% B, 13-20 min from 45% B to 99% B, and 20-24 min held at 99% B. The analytical solvent gradient was followed by a 16 min re-equilibration gradient to prep the column before the next sample injection that went as follows: 0-0.05 min held at 99% B at 0.25 mL/min, 0.05-1 min from 99% B to 50% B and 0.25 mL/min to 0.1 mL/min, 1-11 min held at 50% B and 0.1 mL/min, 11-11.05 min from 50% B to 0% B at 0.1 mL/min, 11.05-14 min held at 0% B at 0.1 mL/min, 14-14.05 min held at 0% B and increased flow rate from 0.1 mL/min to 0.25 mL/min, and 14.05-16 min held at 0% B and 0.25 mL/min. Data were collected on an Orbitrap Exploris 240 using a heated electrospray ionization (H-ESI) source in ESI negative mode. The mass spectrometer acquisition settings were as follows: source voltage -2,500V, sheath gas 60, aux gas 19, sweep gas 1, ion transfer tube temperature 320ºC, and vaporizer temperature 250ºC. Full scan data were collected with a scan range of 70-800 m/z at a mass resolution of 240,000. Fragmentation data was collected using a data-dependent MS2 (ddMS2) acquisition method with MS1 mass resolution at 120,000, MS2 mass resolution at 15,000, and HCD collision energy fixed at 30%.
